# Characterizing Locus Specific Chromatin Structure and Dynamics with Correlative Conventional and Super Resolution imaging in living cells

**DOI:** 10.1101/2021.03.16.435731

**Authors:** Dushyant Mehra, Santosh Adhikari, Chiranjib Banerjee, Elias M. Puchner

## Abstract

The dynamic rearrangement of chromatin is critical for gene regulation, but mapping both the spatial organization of chromatin and its dynamics remains a challenge. Many structural conformations are too small to be resolved via conventional fluorescence microscopy and the long acquisition time of super-resolution PALM imaging precludes the structural characterization of chromatin below the optical diffraction limit in living cells due to chromatin motion. Here we develop a correlative conventional fluorescence and PALM imaging approach to quantitatively map time-averaged chromatin structure and dynamics below the optical diffraction limit in living cells. By assigning localizations to a locus as it moves, we reliably discriminate between bound and searching dCas9 molecules, whose mobility overlap. Our approach accounts for changes in DNA mobility and relates local chromatin motion to larger scale domain movement. In our experimental system, we show that compacted telomeres have a higher density of bound dCas9 molecules, but the relative motion of those molecules is more restricted than in less compacted telomeres. Correlative conventional and PALM imaging therefore improves the ability to analyze the mobility and time-averaged nanoscopic structural features of locus specific chromatin with single molecule precision and yields unprecedented insights across length and time scales.

## Introduction

The spatio-temporal organization of chromatin regulates gene expression on various length and timescales. On the smallest length scale of individual nucleosomes (~10 nm), which is also referred to as the primary structure of chromatin, covalent modification of histone tails regulates the interaction of nucleosomes with DNA and therefore chromatin accessibility^1–3^. The secondary structure integrates features of the primary structure in organizations up to ~100 nm^1–3^. The tertiary structure of chromatin comprises larger domains such as enhancer promoter contacts to regulate gene expression^1,3^. In addition the regulation of DNA accessibility controls transcription factor binding^4^. Even larger chromatin structures such as topologically associated domains can create long range contacts for instance in super-enhancer complexes to activate transcription^2^. Many sequencing based methods have been used to indirectly characterize chromatin structure across these length scale^4^. However, these methods only provide time-averaged structural information of a population of cells or a structural snapshot of a single cell^2,5^. Recent evidence suggests that chromatin compaction, condensation, and accessibility is dynamically regulated at both small and large length scales as well as fast and slow time scales^6^. The degree in correlated movement between small chromatin structures and the large chromatin domains they reside in has been recognized as an important feature of nuclear phase condensates, which are hypothesized to help regulate gene expression^6–11^. Therefore, studying the nanoscopic structure and dynamics of chromatin across time and length scales is needed to characterize how genome organization affects gene regulation^6,7,12^. Here, we present a correlative conventional fluorescence and photo activation localization microscopy (PALM) approach that overcomes current limitations to study the relation between chromatin structure and dynamics at specific loci across various time and length scales in living cells.

Recently, CRISPR/dCas9-based fluorescence imaging methods have been developed as a modular approach to map chromatin structure and dynamics. By using programmable guide RNAs (gRNA), fluorophores can be targeted to specific sequences in the genome^13–15^. These methods require tens of fluorescent probes to be bound to a locus of interest in order to overcome the background fluorescence from the majority of freely diffusing and searching probes^13,15–17^.To reduce the number of gRNAs needed to detect the fluorescence signal of a locus, repetitive RNA aptamers such as MS2 sequences have been appended to gRNA^18–21^. By employing these labeling strategies in conjunction with conventional time lapse fluorescence imaging, valuable information about the slow and long-term dynamics of loci has been obtained. However, the optical diffraction limit precludes the ability of conventional fluorescence imaging to map the organization of smaller chromatin structures below ~250 nm. With the advent of photoactivated localization microscopy (PALM), it became possible to track single molecules in living cells and to resolve intracellular structures in fixed cells with ~20 nm resolution^22,23^. By employing sparse activation of photoactivatable or photoswitchable fluorophores, PALM avoids spatio-temporal overlap of dense emitters and allows for the determination of the precise locations of individual fluorophores by Gaussian fitting. Over time many localizations are then accumulated for linking them to molecular trajectories or for resolving structures below the optical diffraction limit. Recently, CRISPR/dCas9-based DNA labeling has been combined with PALM to monitor chromatin dynamics at the nucleosomal level in living eukaryotic and prokaryotic cells or to obtain structural information such as chromatin compaction or condensation in chemically fixed cells^24–30^. However, the long acquisition times required for PALM imaging preclude the ability to obtain both, structural and dynamic information in living cells because the motion of DNA spreads out the localizations of fluorophores bound to a locus along its trajectory. Therefore, it is a challenge to relate chromatin structure and dynamics across length and time scales and to characterize dynamic chromatin rearrangements below the optical diffraction limit^6^.

Correlative single molecule and conventional (or stimulated emission depletion) approaches have been previously used to study the nanoscale organization and the dynamics of various biological structures in living cells^31–37^. These studies involve dual tagging of the same protein or of interacting proteins with a conventional and a PALM compatible fluorophore. In this way, it has been possible to observe how the protein appended with the PALM tag moves relative to the conventionally tagged protein. These methods have been employed to characterize protein interaction kinetics and to obtain both dynamic and structural information about specific organelles^31–37^. However, currently there is no method in living cells to obtain super resolved structural information of specific loci while they move, to relate structural and dynamic information at a locus, or to compare small with large scale motion at a locus.

Here we develop a correlative conventional fluorescence and PALM imaging approach in living cells to bridge these gaps of current techniques. This approach allows to resolve and quantify locus specific chromatin structures by correcting for their motion. In addition, single fluorescent probes are more reliably identified to be associated with a moving locus in order to determine their mobility state. We demonstrate correlative conventional and PALM imaging using the well characterized telomere sequences as a model system. First, we tagged dCas9 and the MS2 coat proteins (MCP) that bind to a modified telomere-targeting gRNA scaffold with a conventional and a spectrally distinct PALM fluorophore. Next, each telomere is tracked with the conventional fluorescence signal while the single molecule localizations are recorded in the PALM channel of the microscope. The trajectory of the locus is then subtracted from the coordinates of the single molecule localizations to correct for motion blurring. As a result, structural parameters such as the time-averaged size of a locus or the density of bound probes can be quantified, which can yield insights into the compaction of chromatin. By determining the location and mobility of a locus relative to the traces of single dCas9/MCP complexes during imaging, dCas9/MCP complexes can be more reliably identified to be bound to a locus. This relative motion of single molecules compared to the locus shows how smaller scale nucleosomal rearrangements contribute to larger scale chromatin structural movements. Furthermore, we relate the compaction and condensation of telomere clusters to their local and global chromatin mobility. This study demonstrates that correlative conventional fluorescence and PALM imaging accurately identifies Cas9 molecules bound to a locus and yields quantitative dynamic and structural information about specific genomic loci at the nanoscale in living cells.

## Results and Discussion

### Motion of telomeres prevents high-resolution PALM measurements in living cells

In order to demonstrate the challenges of conventional PALM imaging to assign localizations to a locus and to extract structural information, we targeted the highly repetitive telomere sequence that has been previously characterized with conventional fluorescence and PALM imaging^13–15,26,28^. We transiently expressed a GFP-tagged dCas9 for conventional fluorescence imaging in addition to the MCP-HaloTag protein, which was conjugated to the PALM compatible PA-JF646 dye and binds to the gRNA MS2 sites and forms a complex with dCas9-GFP. In the absence of gRNA, both the conventional fluorescence signal of dCas9-GFP and the single molecule localization from MCP-HaloTag were distributed throughout the nucleus (Fig. 1A). This result is expected since both fluorescent reporters cannot bind to the telomere regions in the absence of gRNA. Importantly, the single molecule localizations exhibited some degree of clustering, which highlights the difficulty to identify which localizations are bound to a locus just based on their density. When the gRNA was expressed, the conventional fluorescence signal showed pronounced puncta indicating dense binding of dCas9-GFP to telomeres as in previous studies^13,15,28^. The single molecule localizations in the PALM images also formed dense regions that partially overlapped with the conventional fluorescence puncta. However, the single molecule localizations were smeared out to various lengths along the movement of the telomeres during the long PALM data acquisition time. This result demonstrates that high-resolution structural information cannot be obtained via PALM imaging in live cells and that the motion of loci needs to be corrected.

**Figure 1:**
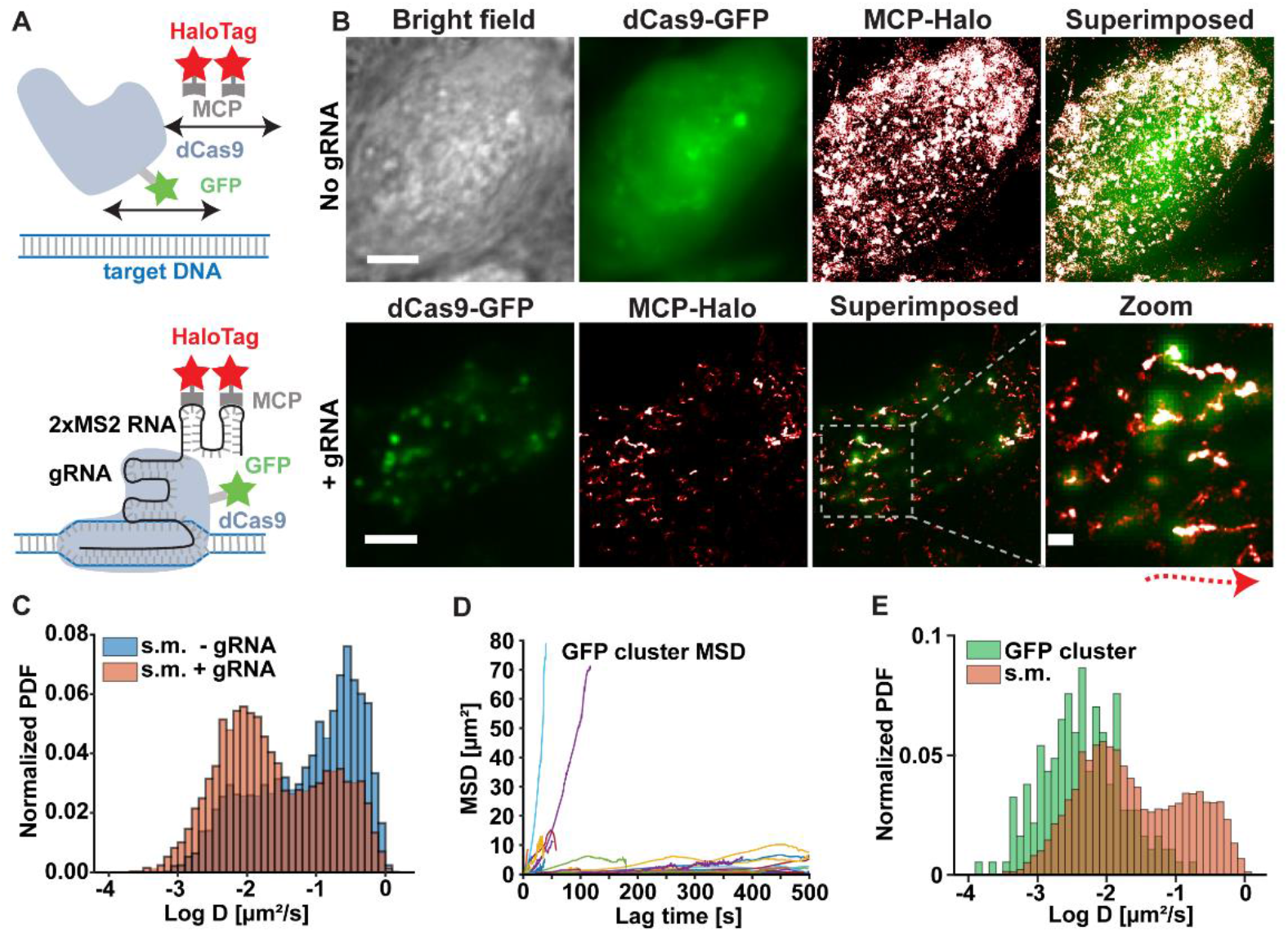
Motion of telomeres prevents high-resolution PALM measurements in living cells. **A)** Fluorescent labeling strategy of repetitive telomere sequences involving GFP-tagged dCas9 for conventional fluorescence imaging and MCP-HaloTag proteins for PALM imaging that bind to the two MS2 sites of the guide RNA and form a complex with dCas9 (lower). B) Upper: In the absence of gRNA the conventional fluorescence signal of dCas9-GFP (green) as well as the PALM localizations of MCP-HaloTag (red/white) are distributed throughout the nucleus of a cell. Lower: In the presence of gRNA dCas9-GFP and MCP-HaloTag bind to and form a complex at the telomere sites as evident by the presence of bright puncta in the GFP and PALM images. The magnified superposition shows clear telomere clusters in the GFP image while the PALM localizations are smeared out at various degrees along the paths of the telomeres during the long PALM data acquisition time. Scale bar represents 5 μm. Scale bar in zoom image represents 800 nm. C) The diffusion coefficient distribution from single MCP-HaloTag proteins significantly overlaps in cells with and without telomere gRNA (7281 traces with telomere gRNA and n = 5749 traces without gRNA from N = 5 cells). D) The mean-square displacement of telomere GFP traces exhibits a wide variability with some immobile telomeres and others that are actively transported (n = 81 traces from N = 5 cells). E) The diffusion coefficient distributions of GFP telomere traces and single MCP-HaloTag traces show some degree of overlap (N = 5 cells).

With the same PALM data we performed single molecule tracking of MCP-HaloTag by linking localizations that appeared within a distance threshold in consecutive frames (see materials and methods). From the resulting single molecule traces, the mean-square displacements (MSDs) and corresponding diffusion coefficients were calculated. In the presence of gRNA, the distribution of MCP-HaloTag proteins exhibited a pronounced peak at large diffusion coefficients around 0.22 +/− 0.01 μm^2^/s, indicating fast diffusion of searching or unbound probes. In addition, the diffusion coefficient distribution contained a peak at small diffusion coefficients around 0.009 +/− 0.003 μm^2^/s, which could be interpreted as the mobility of bound probes (Fig. 1C). The distribution of diffusion coefficients in the absence of gRNA also showed a peak at large diffusion coefficients (0.28 +/− 0.03 μm^2^/s) but in addition contained a significant fraction of small diffusion coefficients. Importantly, these small diffusion coefficients overlapped with the slow fraction of bound MCP proteins in the presence of gRNA. The overall overlap between both distributions of 71.5% demonstrates that single molecule traces from the dCas9/MCP complex cannot be classified as bound just by their diffusion coefficient. This issue is further compounded by the increased uncertainty of diffusion coefficients obtained by fitting short traces due to increased effect of the localization error. It is important to note that the diffusion coefficient distribution of a fluorophore with a nuclear retention signal shows the same slow diffusing fraction as MCP in the absence of gRNA. (Sup. Fig. 2C). The Kolmogorov Smirnov test performed at a 0.05 significance level showed that the diffusion coefficient distributions are not statistically different (P = .79). Therefore, the slow fraction of MCP proteins in the absence of gRNA is not caused by non-specific protein RNA interactions but is likely due to the inhomogeneous and highly crowed environment of the nucleus that can have a wide range of apparent viscosities^24,28–30,38,39^.

In order to quantify the mobility of telomeres and to determine the extent of motion blurring in PALM images, we tracked the conventional fluorescence signal that was recorded every 10th frame of the PALM imaging sequence. The resulting mean-square displacement vs. time traces exhibited a very wide range of slopes with some telomeres being almost immobile, while others were actively transported over a distance of up to 10 μm (Fig. 1D). This heterogeneity of telomere mobilities highlights the need to correct PALM localizations of each individual telomere for motion in order to obtain structural information. Furthermore, the diffusion coefficient distribution of telomere traces exhibited partial overlap with the bound fraction of single MCP traces (Fig. 1E).

However, single molecule traces are generally faster, which can be caused by a relative motion with respect to the center of mass of telomeres and the additional underlying motion of the telomeres themselves. The heterogeneity of the diffusion coefficients of individual telomeres will therefore cause errors in quantifying the mobility of bound MCP molecules and in identifying bound MCP molecules based on a mobility threshold (Fig. 1D and 1E).

In summary, the differences in chromatin mobility and the overlapping diffusion coefficient distributions of bound and unbound dCas9/MCP complexes result in errors for identifying bound single molecules based on their mobility and for quantifying their mobility. Furthermore, the wide range of telomere mobilities results in significant spreading of their bound single molecule localizations, preventing the extraction of structural parameters. These results demonstrate the need to employ a correlative conventional fluorescence and PALM imaging approach to dynamically identify these bound molecules throughout the long PALM data acquisition time and to correct for motion.

### Correlative conventional fluorescence and PALM imaging identifies bound molecules and corrects for chromatin motion

In order to correct for telomere motion during the long PALM data acquisition time and to reliably identify bound single molecules, we developed a correlative conventional and PALM imaging approach. In this approach telomeres are labeled with a conventional fluorophore such as GFP to continuously track their trajectory throughout the entire data acquisition time. In addition, telomeres are labelled with a spectrally distinct PALM dye (PA-JF646 bound to MCP HaloTag) to localize sparse single molecule signals, which get smeared out along the trajectory of a telomere (Fig. 1B). By subtracting the trajectory of the telomere from the single molecule coordinates, the motion can be corrected and a super-resolution images can be recovered and further quantified.

To minimize bleaching of dCas9-GFP, we employed a repetitive 10 frame shutter sequence, in which GFP is excited in one frame followed by photoactivation in the next frame and 646 nm excitation in the remaining 8 frames. As seen in Fig. 2B, telomeres were detected and localized as continuously fluorescing puncta during the GFP excitation frames, whereas sparse and bright fluorescence bursts from single molecules were detected during the remaining PALM frames. These bursts were then localized by gaussian fitting as in conventional PALM data analysis and rendered as gaussians with a width equal to the localization precision. The resulting PALM images exhibited single molecule localizations throughout the nucleus with some clustered regions and dense tracks along the motion of telomeres (Fig. 2C. left). However, the effect of motion blurring and the background localizations of unbound molecules prevent the extraction of structural parameters and the reliable identification of molecules bound to telomeres. In order to assign single molecule localizations to a telomere, the GFP localizations were linked to trajectories of individual telomeres and linearly interpolated during the PALM frames (Materials and Methods). The interpolation over 10 frames results in and estimated median error of 45 +/− 10 nm (Sup. Fig. 3A and 3B). Single molecule localizations were then assigned to the nearest telomere GFP localization if they were separated by less than the radius of the GFP cluster. In addition, the single molecule localizations were linked to traces (Materials and Methods) to further exclude MCP molecules in their freely diffusing state or dCas9/MCP complexes in their search state on the DNA. If a molecule entered a telomere but remained bound for at least four frames, it was included. If a molecule entered and exited a telomere, it was excluded and interpreted as a searching or freely diffusing molecule (see Materials and Methods). This assignment of traces is based on previous studies showing that the residence time of bound Cas9, the stability of gRNA and the residence time of MCP bound to the gRNA is on the order of minutes to hours^20,24,29,40,41^. On the other hand, dCas9 residence time during scanning has been measured to be 20-30 milliseconds in prokaryotic genomes and between 20-100 ms in eukaryotic genomes^25,27,29^. Based on these experimentally determined upper and lower time limits and the uncertainty for calculating the mobility for shorter traces, a 4 frame cutoff was chosen as a compromise to accurately determine the mobility state of a protein and to not falsely assign a scanning protein to be bound to the locus.

**Figure 2:**
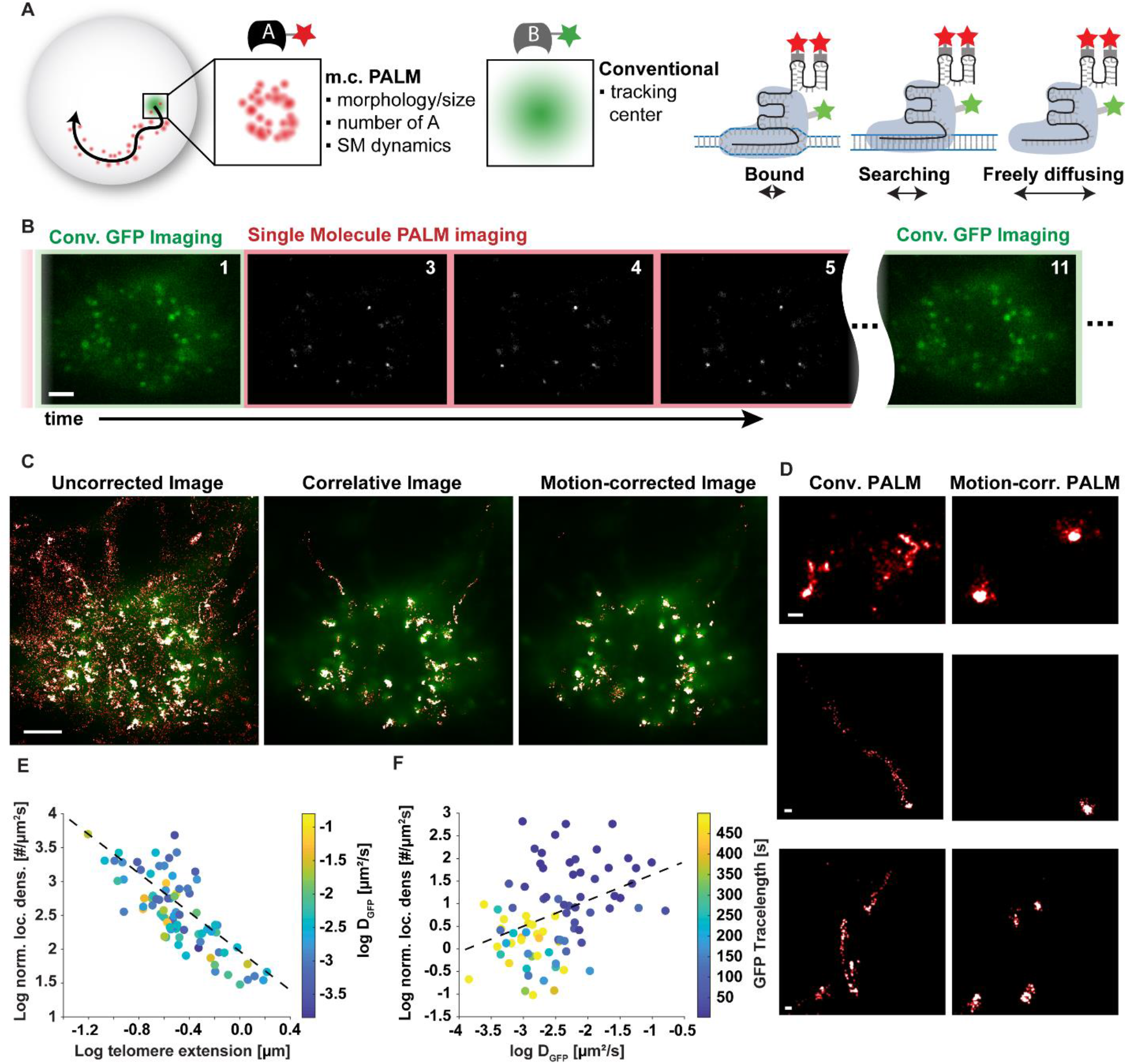
Correlative conventional fluorescence and PALM imaging identifies bound molecules and super-resolves moving chromatin. A) Schematics of motion correction via correlative conventional and PALM imaging. Left: Single molecule localizations are smeared out along the trajectory of a locus, which is simultaneously tracked in the GFP channel. By subtracting the trajectory from PALM localizations the moving locus can be super-resolved. Right: In addition, correlative imaging can better assign bound localizations at any instance in time by the proximity to the conventional GFP signal and reduces background from searching and freely diffusing probes. B) In a correlative conventional and PALM imaging sequence, GFP is excited and imaged in one frame followed by one frame of 405 nm photoactivation and brighfield imaging (not shown) and 8 frames of PALM imaging, in which bright single molecule fluorescent signals become visible. C) Left: Superposition of a GFP image and conventional PALM image that includes a majority of freely diffusing and searching fluorescent probes. Middle: The correlative conventional and PALM image only depicts PALM localizations that appear in proximity to a GFP cluster at any instance in time and suppresses background from freely diffusing and searching probes. Right: The motion-corrected PALM image super-resolves each moving telomere, which co-localizes with its GFP signal. More telomeres are visible in the GFP channel since they are detected over a wider z-range compared to PALM. Scale bar represents 5μm D) Magnifications of correlative PALM and motion-corrected PALM images show localizations of telomeres smeared out to various degrees (left) and super-resolved telomeres (right). Scale bar in all images in panel D represent 200 nm. E) The localization density of individual telomeres is obtained by motion-correction PALM and shows a negative correlation with the maximum extension of telomeres (Correlation Coefficient = - 0.77). The color of each telomere indicates no clear dependence on the GFP diffusion coefficient. F) The localization density of telomeres shows a slight correlation (Correlation Coefficient = 0.29) with their diffusion coefficient. The color represents the temporal tracelength of the GFP telomere signal and indicates that slower telomeres can be imaged for a longer time. All data taken is from n = 81 clusters across N = 5 cells.

In the resulting PALM image depicting only localizations that were bound to a telomere, most background localizations are suppressed, and motion-blurred telomeres become clearly visible (Fig. 2C middle). After subtracting the GFP-trajectories of individual telomeres from their single molecule localizations, motion blurring is corrected, and a super-resolution image of each telomere is obtained (Fig. 2C right). The individual examples shown in Fig. 2D highlight how this method is able to correct for the significant motion of a telomere and to recover the time-averaged super resolved structure of a locus.

The motion-corrected super resolution image of each telomere provides quantitative structural information such as the degree of chromatin compaction and condensation. The area of the motion corrected cluster, for instance, quantifies the size of each telomere averaged over the time of data acquisition. The time-averaged size measurements of telomeres obtained with our method are consistent with previous measurements in chemically fixed cells^13,26,28,42^. However, it is important to note that differences in telomere size can be due to differences in telomere lengths in different cell types, different chromatin states, or expansion or compaction of loci during live cell imaging^13,28,43^. This information can then be related to other quantities such as the number of localizations bound to a locus to obtain insights into the density of bound probes (Fig 2E). We observed an overall trend of smaller, less extended telomeres having a higher localization density whereas larger and more extended telomeres exhibited a lower localization density (Fig. 2E). Since the maximum extension of telomeres depends to some degree on their area this observation can be explained by less extended telomeres being more compact and exhibiting a higher density of bound fluorophores while more extended telomeres having a lower fluorophore density (Sup. Fig. 4C).

While localization density measurements of chromatin have been performed in fixed cells in the past and used as a measure of chromatin condensation^26,28,42,44^, the power of motion-correction PALM is to perform such measurements in living cells and to simultaneously obtain dynamic information about the motion of individual telomeres and of individual bound fluorescent probes. We therefore related the localization density of each telomere to their mobility determined from their GFP signal (Fig. 2F). The mobility of telomeres did not show a tight correlation with their localization density (Correlation Coefficient = 0.39), however, slowly moving telomeres generally had a lower localization density while faster telomeres had a higher localization density (Fig, 2F). This observation can be explained by the ability of dense telomeres that are generally also less extended to move more freely while less dense and larger telomeres might be more restricted in their movement. This is consistent with previous studies comparing indirect telomere size measurements and mobility^13^. Variations in telomere mobility across similar sizes and densities could be explained by local changes in the chromatin environment (Fig 2F)^13^. Importantly, since the activation rate for MCP-HaloTag localizations was held constant during imaging, the number of localizations has been normalized by the observation time/GFP temporal tracelength of telomeres (Sup. Fig. 4A). Localization densities of telomere clusters of varying temporal trace lengths can therefore still be compared without bias. (Sup. Fig. 4A, 4B). However, telomeres with a large diffusion coefficient were observed a shorter time than slow telomeres since faster telomeres are more likely to move out of focus during the data acquisition time (Fig. 2F).

These results demonstrate that motion correction PALM is able to provide quantitative time averaged structural insights e.g. into the compaction and condensation across time and length scales in living cells and to relate this structural information to the dynamics of loci.

### Reliable identification of single molecule traces that are bound to moving loci

Correlative conventional and PALM imaging not only yields structural information of chromatin as shown above, but can also be applied to study how DNA at the nucleosomal level moves relative to the larger chromatin domain it resides in. Recent studies highlight the importance of characterizing the degree of correlation between small- and large scale chromatin motion and its relation to gene regulation and phase separation^6–8,10,45^. A prerequisite for measuring relative DNA mobility is the ability to reliably identify dCas9/MCP complexes that are bound to a locus and to separate them from searching and freely diffusing ones. Since the diffusion coefficient distributions of these three mobility states overlap (Fig. 1C), it is not possible to identify bound Cas9/MCP complexes just based on their mobility. Here, we demonstrate that correlative conventional and PALM imaging reliably identifies traces of single dCas9/MCP complexes bound to chromatin in order to better characterize their mobility without imposing any threshold on diffusion coefficients.

Using correlative conventional and PALM imaging, dCas9/MCP traces were classified as bound, unbound and partially bound based on their proximity to a conventional fluorescence signal from a telomere at each point in time as described in the previous section (see also Materials and Methods). The diffusion coefficient distributions of bound and partially bound trace were unimodal with one peak at 0.007 +/− 0.001 μm^2^/s and 0.018 +/− 0.002 μm^2^/s respectively, while the unbound trace distribution exhibited two distinct peaks at 0.01 μm^2^/s +/− 0.003 and 0.22 +/− 0.005 μm^2^/s (Figure 3A). The diffusion coefficient distribution of unbound traces had significant overlap with the one of bound traces, highlighting the inability to classify bound dCas9/MCP complexes just based on their mobility. For instance, the normalized probability density distribution of bound and unbound traces had a 62% overlap (Sup. Fig. 5A and 5B). Correlative conventional and PALM, however, is able to reliably separate slowly moving unbound traces from bound dCas9/MCP complexes. These results demonstrate that an accurate classification of dCas9/MCP complexes can be used to study their dynamics while freely diffusing, searching on DNA, and while being bound to a target locus.

The advantages of accurately classifying bound traces also manifest themselves when comparing the motion of individual dCas9/MCP complexes to the motion of the entire telomere they reside in. The diffusion coefficient distributions of all dCas9/MCP traces only exhibit a partial 68.5% overlap with the ones of the telomere cluster traces (Fig. 1E). However, the traces classified with correlative imaging to be bound show a significantly larger overlap of 75% and a high degree of correlated motion (Fig. 3C, Sup. Fig 5C and 5D). Both diffusion coefficient distributions resemble a Gaussian distribution on a log scale and indicate sub-diffusive mobility, which is a hallmark of chromatin mobility. While bound dCas9/MCP traces and entire telomeres exhibit sub-diffusive mobility, the median of the diffusion coefficient distribution of the bound traces (0.007 +/− 0.003 μm^2^/s) is larger than the one of telomere cluster traces (0.002 +/− 0.0005 μm^2^/s). We hypothesize that this difference is due to the fact that DNA at the nucleosomal level tends to move faster than its larger domain counterpart. In contrast, the skewness in the diffusion coefficient distribution of partially bound traces compared to bound dCas9/MCP and telomere traces indicates that the mobility of these molecules is not sub-diffusive. Therefore, partially bound molecules are not likely bound to DNA but likely diffusing through a telomere cluster or searching for a binding site. This interpretation is further supported by previously reported residence times of dCas9 on DNA on the order of minutes to hours, which is much longer than the bleaching time of fluorescent probes used for localization microscopy^24^. In addition, nucleosomes have been shown to kick off scanning dCas9 molecules and shorten search state residence time to milliseconds in eukaryotic cells^24^. Because of these two facts, it is very unlikely to observe a dCas9/MCP complex bind to and leave its target site during the length of a single molecule trace.

**Figure 3:**
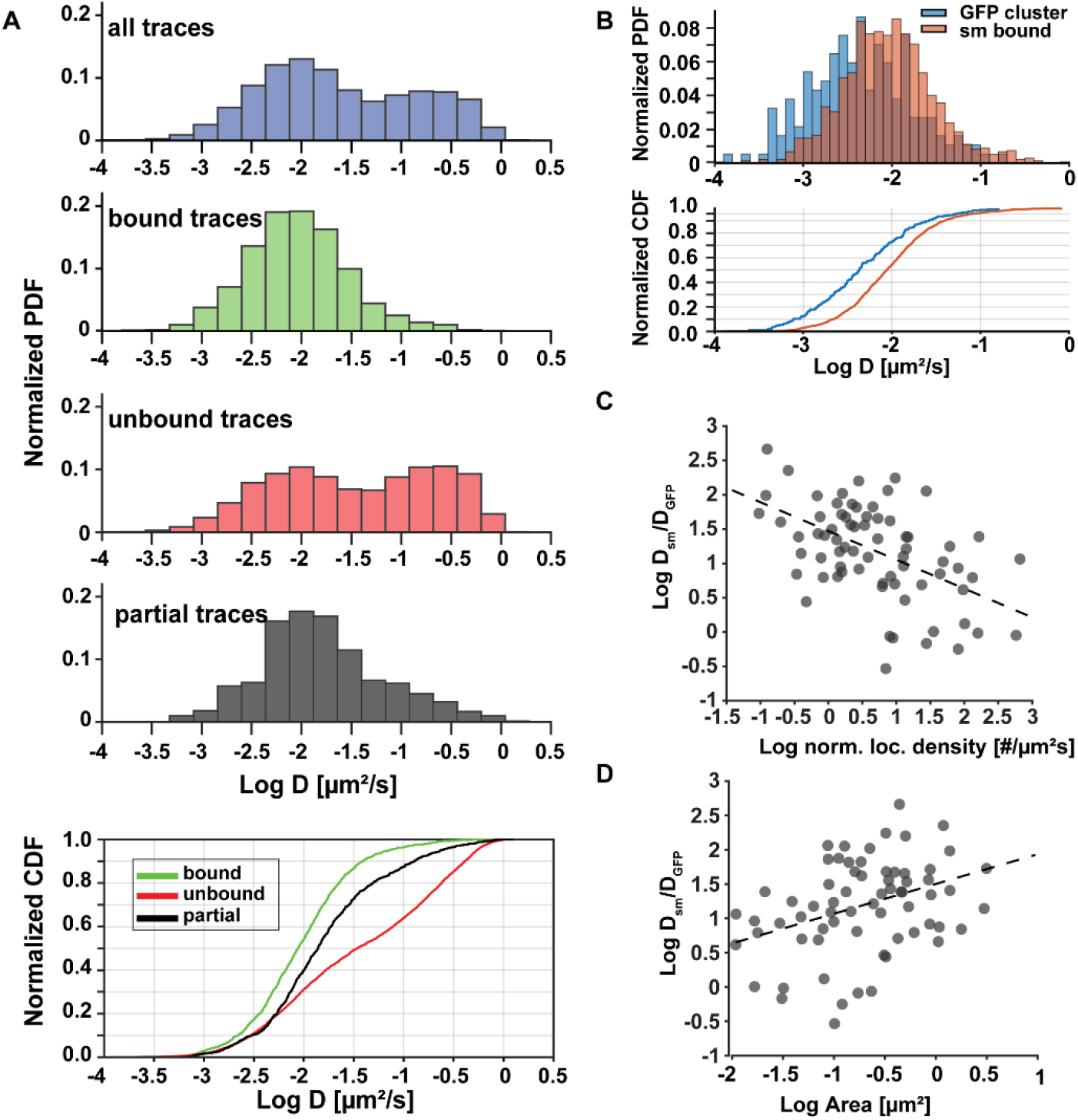
Accurate identification of bound dCas9 molecules reveals their telomere-specific relative mobilities. A) Upper: Diffusion coefficient histograms of all dCas9-MCP complexes and of molecules identified with correlative conventional and PALM imaging to be bound (n = 1490 traces), unbound (n = 5077 traces) and partially bound (n = 714 traces) to telomeres. Lower: Cumulative diffusion coefficient histograms of the same classes of molecules. There is significant overlap of all distributions. B) Upper: Diffusion coefficient histogram of telomeres (GFP) and dCas9-MCP complexes identified to be bound. There is significant overlap but generally single molecules move faster than the telomeres they reside in. Lower: Cumulative diffusion coefficient histogram of the same classes. C) The ratio of average dCas9-MCP diffusion coefficients and the telomere they reside in shows a negative correlation (Correlation Coefficient = - 0.52) with the normalized localization density of the telomeres. D) The ratio of average dCas9-MCP diffusion coefficients show a positive correlation (Correlation Coefficient = 0.48) with the area of the telomeres they reside in. All data was collected from N = 5 cells and there were no statistically significant differences in the group comparisons between cells (multi-way ANOVA p=0.35).

We next evaluated how fast single dCas9/MCP complexes move relative to the larger telomere domain they reside in. The average diffusion coefficient of all bound traces in a telomere cluster was calculated by fitting the averaged MSDs to the 2D diffusion equation and then divided by the diffusion coefficient of the telomere (Fig. 3C and 3D). In almost all cases, single dCas9/MCP complexes move faster than their larger telomere domain they reside in (Fig. 3C and 3D). In a few cases the single molecule traces were coincidentally obtained during sections where the mobility of the cluster was faster than the average mobility of the telomere cluster. Telomeres with a low localization density exhibited the fastest relative mobility of single dCas9/MCP complexes and the largest variability in their relative mobilities (Sup. Fig. 5E). dCas9/MCP complexes in dense telomeres had a low or no relative mobility and a more narrow range of diffusion coefficient ratios (Fig. 3D). These results indicate that less dense telomeres undergo a high degree of dynamic chromatin rearrangement while dense telomeres exhibit a more static chromatin state potentially due to tighter interactions. When comparing the relative diffusion coefficient with the area of telomeres, a slight correlation is observed (Sup. Fig. 5F). Smaller clusters that are potentially more dense showed the lowest relative mobility of single molecules whereas larger clusters showed the highest relative mobilities (Sup. Fig. 5E and 5F). This result is consistent with the previous finding that denser telomeres are smaller (Fig. 2E and Sup. Fig. 4C) and again indicates that larger, less compacted telomeres undergo more chromatin remodeling compared to small telomeres (Fig 3D Sup. Fig. 5F.).

In summary, these results demonstrate that correlative conventional and PALM imaging can reveal how chromatin compaction and condensation affects the motion of small nuclear rearrangements within a larger chromatin domain. Overall, this data shows that dCas9/MCP complexes move significantly faster than the larger telomere domains they reside in. We show that more compact clusters with a higher localization density, exhibit a higher degree in correlation in single molecule mobility and less variance compared to less dense clusters. These findings are consistent with single nucleosome tracking measurements showing that more compact chromatin domains move coherently^46–48^. Our findings also match existing nucleosome tracking data that identified chromatin density as an important regulator of instantaneous chromatin dynamics^6^. Since the relative mobility of chromatin is a hallmark of protein DNA phase condensation and gene regulation, our approach may aide in characterizing the formation of nuclear phase condensates in future experiments with appropriate controls^6–8,10,12,45^.

### Mobility state analysis reveals hidden state in bound traces

A general limitation to obtain accurate diffusion coefficients from short single molecule traces is the increased effect of the localization uncertainty^49,50^. Only an apparent diffusion coefficient can be obtained and used as a relative mobility comparison to other traces. To obtain a more accurate diffusion coefficient estimate, traces or individual step-sizes are therefore often clustered into specific mobility populations and the average diffusion coefficient of the population is calculated^51,52^. We therefore utilized the Bayesian cluster analysis SMAUG to determine mobility states of dCas9/MCP complexes and to demonstrate that correlative conventional and PALM imaging can reveal mobility states that are hidden when all traces of conventional PALM data are analyzed^51^. We used this method over Gaussian mixture models, hidden Markov models, and other vibrational Bayesian clustering techniques due to the ability of the SMAUG algorithm to analyze short traces without the requirement of a predetermined number of mobility states for fitting single molecule data^51^.

First, we analyzed the mobility states of all dCas9/MCP single molecule traces with SMAUG. The result in Fig. 4A (upper left) shows the weight fraction of the mobility state classification and the apparent diffusion coefficient of each mobility state from the last 20,000 iterations of the SMAUG algorithm (see also Sup. Fig. 6A,B,C,D). SMAUG identified a bound and faster unbound state with clear convergence. However, it is important to note that the absolute values of diffusion coefficients obtained by SMAUG do not match the ones obtained by mean-square displacement fitting due to its algorithm that takes into account localization uncertainties and only analyzes displacements of single molecules in consecutive frames. Therefore, only the relative classification of traces into different mobility states was used to then calculate the diffusion coefficient distribution based on MSD fitting (Fig. 4A, lower). The traces classified by SMAUG to contain bound step sizes exhibit a similar distribution as the ones obtained from the correlative approach with a peak at .01 μm^2^/s (Fig. 3A). Likewise, the traces classified as unbound exhibited a bimodal distribution with an additional peak at 0.5 μm^2^/s. This result is consistent with the control experiment in Fig. 1C in the absence of gRNA, where a slow diffusing unbound population of dCas9/MCP was observed. These two peaks in the diffusion coefficient distribution are not visible in the overall assignment of mobility states by SMAUG (Fig. 4A, upper) since its algorithm uses displacements in consecutive frames and assigns an averaged diffusion coefficient to the population. Therefore, a trace containing a step of the unbound population could have other shorter steps that result in a slow diffusion coefficient from its MSD.

**Figure 4:**
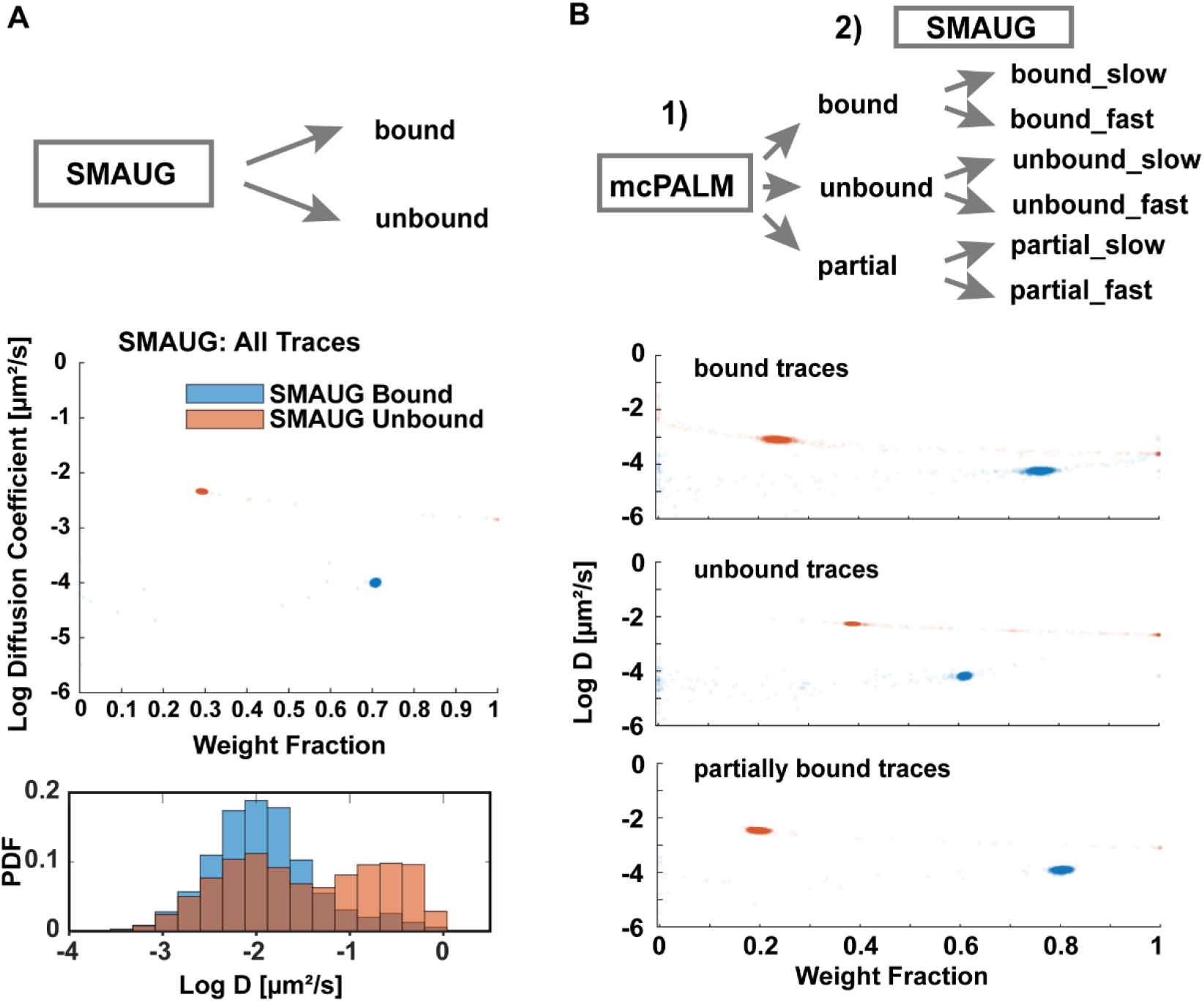
Correlative conventional and PALM imaging combined with SMAUG analysis reveals hidden mobility states. A) Upper: When all dCas9-MCP traces are analyzed with SMAUG, a bound and unbound population is identified. Lower: calculating the diffusion coefficient distributions from the corresponding bound and unbound traces results in similar distributions as observed with correlative conventional and PALM imaging. B) When correlative conventional and PALM imaging is performed first to separate bound, unbound and partially bound traces, SMAUG identifies in each class two mobility states that were not detected when all traces were analyzed. Data collected from N=5 cells.

We next applied SMAUG to the three classes of traces identified with correlative conventional and PALM imaging to obtain a more refined characterization of mobility states in each class. When the algorithm was applied to the traces identified by motion correction to be bound (Figure 4B), SMAUG identified 2 mobility states. This result could be explained by the heterogeneity in telomere mobility itself and is consistent with other dCas9 and nucleosome imaging data (Figure 1D and 1E)^13,15,28,48^. In both the unbound and partially bound trace populations, SMAUG again identified two mobility states. The fast mobility state identified in the bound population is distinct from the fast and slow mobility states identified by SMAUG in all traces, unbound, and partially bound traces. This shows that when paired with motion correction PALM, SMAUG is able to identify a fast and slow bound population that could not be identified otherwise due to the large overlap in mobilities between slowly moving unbound traces and the bound traces. These results demonstrate that correlative conventional and PALM imaging when combined with other trace classification approaches enhances mobility state analysis by a more accurate identification of the bound population, which can also lead to a more accurate calculation of kinetic parameters.

## Conclusions and Future Outlook

The advent of PALM imaging presented new opportunities to study the structure and dynamics of chromatin in a new level of detail. Previous studies and technique developments yielded new insights into the nanoscopic chromatin structure in fixed cells and into its dynamics in living cells. However, available techniques cannot obtain both, structural and dynamic information in living cells due to the motion of chromatin during the long PALM data acquisition times. Here we developed and demonstrated a correlative conventional fluorescence and PALM imaging approach that tracks the motion of labelled loci in order to correct for their motion in the simultaneously acquired PALM images. We employed dCas9/MCP-based imaging of telomeres as a model system to demonstrate that motion-correction PALM allows for measuring both, time averaged super resolved structural parameters as well as dynamics of chromatin organization such as various transport modes of DNA. This approach also yielded information into how individual dCas9/MCP complexes bound to DNA move relative to larger chromatin domains they reside in. Identifying bound dCas9/MCP probes using the correlative approach avoids the need of using a fixed mobility threshold and provides more accurate information about single molecule probes bound to chromatin.

Our study lays the foundation for further refinement and optimization of the presented correlative conventional fluorescence and PALM imaging technique and its application to study a myriad of different chromatin loci. For instance, extending this technique to 3D to track loci for longer periods of time and to extend the lifetime of the conventional signal would extend the imaging time of loci and result in an improved spatial and temporal resolution^15,28,44,53,54^. In addition, orthogonal RNA binding proteins and/or CRISPR-Cas proteins could be applied to enable simultaneous multicolor imaging of different chromatin regions^14,18–20^. Because dCas9 can only bind to accessible chromatin regions, characterizing the number of bound probes at a specific locus can shed light into the chromatin accessibility and show how it changes over time or in response to perturbations^40,55,56^. Correlative conventional and PALM imaging therefore paves the way to study chromatin structure and dynamics of locally repetitive or non-repetitive loci in a new level of detail and to relate the obtained structural and dynamic parameters in living cells.

## Materials and Methods

### Plasmid generation

2XMS2 gRNA scaffold and MCP-HaloTag plasmids were obtained from Thoru Pederson through Addgene^19,20^. The CMV-S.p.dCas9-VP64 plasmid obtained from Charles Gerbasch through Addgene was used as the backbone vector. A PCR was performed with eGFP using Thermofisher Platinum Taq High Fidelity to add BsteII and AflII restriction enzyme sites on the N-terminus and C-terminus of GFP respectively with the forward primer: AGTCAGCTAGGAGgtgacccaggagctcccaagaaaaagcgcaaggtaggtagttccgtgagcaagggcgaggagcta and reverse primer GCTGATCAGCGGTTTAAACttaagtttacttgtacagctcgtccatgccgag. The purified PCR product and CMV-dCas9-VP64 backbone were digested using NEB BsteII-HF and AflII-HF restriction enzymes, gel purified and ligated using the NEB T4 DNA ligase kit. Ligated plasmids were amplified with DH5α competent cells and purified using Qiagen Miniprep kit. mEos2 (obtained from Davidson Lab) was cloned into a CMV-MCP-YFP plasmid (from Mazhar Adli from Addgene) to replace MCP-YFP^21^. A PCR was performed on mEos2 using Thermofisher Platinum Taq High Fidelity to add an BstXI restriction enzyme site to the N terminus of mEos2 and to add a nuclear localization sequence followed by a stop codon and XbaI restriction enzyme site to C terminus of the mEos2 using the forward primer: ggagacccaagcttatgggctacccctacgacgtgcccgactacgccatgagtgcgattaagccagacatg and reverse primer aacttaggccctctagatgcatgttatacctttctcttcttttttggtcgtctggcattg. The PCR product and CMV-MCP-YFP plasmid were digested with BstXI and XbaI restriction enzymes, gel purified, ligated and amplified as before. CRISPR gRNAs sequences targeting telomeres were obtained from Bo Huang^13^. Repetitive telomere targeting sequences appended with ACCG and AAAC nucleotide overhangs were obtained as individual oligos from IDT with the sequences accgGTTAGGGTTAGGGTTAGGGTTA and aaacTAACCCTAACCCTAACCCTAAC. The oligos were annealed and cloned into 2xMS2 gRNA scaffold plasmid using previously described protocols^19,20^.

### Cell line culture, transfection and sample preparation

All imaging experiments were performed with Gastrointestinal Stromal Tumor Cells (GIST-T1) provided by Tamas Ordog and Yujiro Hayashi. Cells were cultured in Fluorobrite Media supplemented with 10% Fetal Bovine Serum (FBS), 4 mM L-Glutamine, and 1% Pencillin/Streptomycin to eliminate fluorescence from phenol red. Cells were seeded in No 1.5 8-well plates (IBIDI) at a density of 50000 cells/ml two days prior to imaging. 200 ng of telomere gRNA along with 50 ng of MCP HaloTag and 50 ng of dCas9-GFP plasmids were transfected into GIST-T1 cells 15-17 hours prior to imaging using the Lipofectamine 3000 and p300 reagent. After incubation with transfection reagents, cells were washed twice with serum diluted Fluorobrite media with 1% FBS and 1% Pencillin/Streptomycin. Cells were incubated with 100 nM of PA-JF646 HaloTag dye provided by Bo Huang in serum diluted media for 15 minutes. After incubation cells were washed with serum diluted Fluorobrite media 3 times and placed in a 37°C and 5% CO2 incubator for an additional 30 minutes. The washing process was repeated three additional times prior to imaging to remove unbound PA-JF646 dye that is still able to fluoresce. Cells identified as containing gRNA exhibited distinct GFP clusters throughout the nucleus of cell while cells that weren’t transfected with gRNA didn’t contain these distinct nuclear clusters (Fig. 1B).

### Microscope setup, camera calibration and imaging

The microscope setup and camera calibration that was used for imaging was previously described^57–59^. In short, a Nikon T1 Eclipse inverted microscope equipped with a perfect focus system was used for imaging experiments. All movies were recorded on an Andor iXon 897 Ultra DU-897U electron multiplying charge coupled detector (EMCCD) camera, which was cooled to −70°C and set to an amplifying gain of 30. The 4 excitation lasers (405 nm, 488 nm, 561 nm, 640 nm OBIS-CW, Coherent Optics) were aligned, expanded, and focused into the back focal plane of the objective (Nikon CFI 100x 1.49 NA oil immersion) using a variety of dichroic mirrors, beam expanders, and lenses. Laser intensity modulation was controlled digitally by a computer via USB. A quad band dichoric mirror (ZT405/488/561/640rpc; Chroma) was used to separate fluorescence emission from excitation light. Fluorescence emission was further split into the far red and green signal using a dichroic long pass beamsplitter (FF652-Di01; Semrock) and band pass filters FF731/137 (Semrock) for the far red channel and ET525/50(Chroma) for the green channel. Programmable shutter sequences were inputted into an NI-DAQ board to synchronize laser output with camera frame duration. The HAL4000 software (Zhuang lab github https://github.com/ZhuangLab/storm-control) was used for intensity modulation, programmable shutter sequence input, camera settings, and image acquisition.

For imaging, a 10 frame shutter sequence at 20 Hz was employed to obtain conventional, bright-field LED, and single molecule localization images. The first frame was the conventional GFP image obtained by exciting the sample with a 488 nm laser at 1.75 mW (power density ~100 W/cm^2^). During the second frame a bright field LED image was recorded to observe cell morphology and health while a variable 405 nm intensity between 1–251 μW (power density of ~0.06–15 W/cm^2^) was applied to the cell to photoactivate the HaloTag bound PA-JF646 nm dye. During the following 8 PALM imaging frames, single molecule signals were recorded by exciting cells using a 640 nm laser at 17.5 mW (~1 kW/cm^2^). This shutter sequence was repeated for 15,000-30,000 frames depending on cell health. A bead calibration was employed to accurately map single molecule localizations identified in the 640 nm channel to the 488 nm channel to account for spherical aberrations created by the emission path^58,60^.

For mEos2-NLS experiments, a similar imaging protocol was employed except a 561 nm laser was used for excitation instead of 640 nm laser. In the emission path, fluorescence emission between the red and green channels were separated by a dichroic long pass beamsplitter (T562lpxr BS; Chroma) along with bandpass filters: ET525/50 (Chroma) for the green channel and ET595/50 (Chroma) for the red channel.

### Single molecule localization, single molecule tracking, mean squared displacement and diffusion coefficient estimation

The Insight3 software from Xiaowei Zhuang’s group was used to identify single molecule localizations and to fit them with 2D Gaussians with the following parameters: 7×7 pixel ROI, widths between 250-700 nm, and minimum 100 photons. The x- and y coordinates of the localizations, along with the intensity, width, background, frame number and other parameters were obtained for each identified single molecule. For the 60 Hz MCP-HaloTag experiments and 20 Hz mEos2-NLS experiments, localizations were identified with intensity values above 30 and 50 photons respectively with all other parameters staying similar to the parameters above. For each imaging condition, a minimum of 3 different cells taken at different days was used for analysis. Localizations were obtained as a text file from the Insight3 software and all other analysis codes were written in MATLAB 2018b.

Localizations that were within 0.48 μm of each other in consecutive frames were linked together to a trace. A low number of molecules was activated in each frame to prevent false linking (Sup. Fig. 1A, 1B). Traces with a minimum of 4 localizations in consecutive frames were used for cross correlation and single trace fitting analysis. No dark frame was allowed between localizations. Displacements for each time interval in a single trace were averaged to obtain a time averaged mean squared displacement (TAMSD) vs time plot. Each TAMSD vs time step plot was fitted to the 2D diffusion equation <r^2^>=4DΔt+ 2σ^2^, where D is the diffusion coefficient, Δt is the time step, r^2^ is the TAMSD, and σ is the localization precision.

The diffusion coefficient distributions of MCP-HaloTag with and without telomere gRNA (Figure 1C and Sup. Fig 2A) taken at 20 Hz (fastest possible camera rate for dual color imaging) were statistically similar to diffusion coefficient distributions of MCP-HaloTag with and without telomere gRNA taken at 60 Hz (fastest possible camera rate, Sup. Fig. 2B). The statistical similarity of the distributions was quantitatively assessed using the Kologmorov-Smirnov Test (KS test) (P = .72 no gRNA, P = .65 telomere gRNA). The increase in the upper limit of the diffusion coefficient distributions was the only difference between the distributions taken at 20 Hz and 60 Hz. Similar findings were also observed when comparing diffusion coefficient distributions of dCas9 with and without globally repetitive target gRNAs and telomere binding proteins taken at faster frame rates (50-500 Hz)^24,29,30,61,62^. This indicates that diffusion coefficients of MCP-HaloTag taken at 20 Hz frame rates can identify molecules in their different mobility states.

### GFP cluster identification, tracking and diffusion coefficient estimation

Telomere GFP clusters were also identified using Insight3 software. GFP localizations were obtained by fitting GFP clusters with 2D Gaussians and the following parameters: 11×x11 pixel ROI, widths between 250-7000 nm and a minimum of 200 photons. The mean GFP localization width of telomeres across all analyzed cells was 376 +/− 221 nm and the maximum GFP localization width was 4181 nm. X- and y- coordinates of the GFP localization, along with the intensity, width, background, frame number and other parameters were again obtained for each identified localization.

For telomere cluster tracking experiments, GFP localizations that were within 0.48 μm of each other in consecutive conventional imaging frames (every 10 frames) were linked together to form a trace. Traces with a minimum of 5 localizations were used for downstream analysis. The widths of consecutive localizations in a trace had to be within 200 nm of each other in order for the trace to be included in downstream analysis. An axial bead calibration showed that PSFs width deviations of 200 nm corresponded to axial deviations of 450 nm which is similar to the lateral trace linking threshold^63^. A linear interpolation between the x- and y- coordinates of GFP localizations in a trace in consecutive conventional image frames was employed to obtain GFP coordinates during the frames that contained single molecule localizations. In order to estimate interpolation errors, GFP telomere clusters were conventionally imaged for 200 seconds at 20 Hz. In a 10 frame shutter sequence, frames 2-9 were removed and a linear interpolation was performed between frames 1-10. The interpolated position of the telomere between frames 2-9 was compared to the actual position of the telomere cluster identified at frames 2-9 to calculate the error in interpolation. The median interpolation estimation error is 45 +/− 10 nm (Sup. Fig 2A and 2B) and the mean interpolation error is constant up to 20 frames (Sup. Fig. 2B).

Mean squared displacements and diffusion coefficients were calculated using the same procedure described in the single molecule identification and tracking section.

### Spatio-temporal cross-correlation between single molecule localizations and GFP cluster coordinates and Motion Correction PALM

In order to superimpose the single molecule and GFP localizations for cross correlation analysis, single molecule coordinates were transformed from the 640 nm (top) to the 488 nm (bottom) camera channel. A bead calibration was performed to account for spherical aberrations between the top and bottom camera channel. Multi-wavelength excitable beads (TetraSpeck microspheres, Invitrogen T7279) were placed on a slide and imaged at both 488 nm and 640 nm wavelengths. The positions of the beads in both the top and bottom channel were fitted to a 3^rd^ order polynomial function to extract the coordinate transformation matrix between the two channels^58,60^. Coordinates from at least N=5 movies each with at least 10 sparsely distributed beads over the entire field of view were used to construct the transformation matrix. The bead calibration measurements were performed every day before imaging.

Once single molecule coordinates were transformed, a distance matrix between the interpolated GFP localizations and all single molecule localizations in one frame were calculated. The radius of the last GFP localization prior to interpolation was used as the radius of the interpolated GFP cluster coordinates. Single molecule localizations whose distance to a GFP cluster was smaller than the radius of the cluster plus the localization precision of the cluster and the single molecule localizations were identified as co-localized with telomeres and isolated for further downstream analysis.

Only single molecule localizations that were part of trajectories with at least 4 localizations in consecutive time steps were used for further analysis. Single molecule traces where all localizations resided inside a cluster were identified as bound traces. Single molecule traces where some but not all localizations resided inside a cluster were identified as partially bound and unbound traces. Single molecule traces where none of the localizations resided inside a cluster were identified as unbound traces. To correct for motion of telomeres in PALM images, the coordinates of GFP localizations where subtracted from coordinates of the single molecule localization. Once all the single molecule localizations were motion corrected, a convex hull was applied to the motion corrected localizations in a cluster to find the cluster boundary. This boundary was used to calculate the area. The extension of a cluster was determined to be the furthest distance between two motion corrected localizations that reside within one telomere. Due to the constant photo-activation rate, the number of localizations in each cluster was normalized by duration of that cluster in the field of view (Sup. Fig. 4A, 4B). The normalized number of localizations was used to calculate the localization density.

### Mobility State Clustering using SMAUG

The single molecule analysis using unsupervised Gibbs sampling (SMAUG) program written by the Julie Biteen group was employed to cluster single molecule trace steps into distinct mobility states^51^. Each trace category, all traces, bound traces, partially bound traces, and unbound traces were inputted into the program for mobility state analysis. Each condition was analyzed through SMAUG for 50000 and 100000 iterations 3 separate times to ensure that the algorithm converged into same number of states and weight fractions.

## Acknowledgements

The authors would like to acknowledge Angel Mancebo Jr for writing the two channel bead calibration code and helpful discussions. The authors would like to acknowledge the Julie Biteen group for providing access to SMAUG analysis code and Julie Biteen, Laurent Gollfoy, and Josh Karslake for the helpful discussions on SMAUG and single molecule trace analysis. The authors acknowledge Bo Huang for providing the PA-JF646 dye. The authors acknowledge Bo Huang, Geeta Narlikar, and Li-Chun Tu for the helpful discussions on CRISPR imaging and Cas9 chromatin interactions. The authors acknowledge Michael Schlierf for the helpful discussions on single molecule trace analysis. The authors acknowledge Tamas Ordog and Yujiro Hayashi for providing the GIST-T1 cells. The authors would like to acknowledge Stephen C. Ekker and Karl J. Clark for providing lab space, reagents, and guidance on molecular cloning. The authors also acknowledge Thoru Pederson and for depositing the 2xMS2 gRNA and MCP-HaloTag plasmids from CRISPR rainbow and CRISPR Sirius papers on Addgene, Michael Davidson for depositing mEos2 plasmid on Addgene, Mazhar Adli for depositing the MCP-YFP plasmid on Addgene, and Charles Gerbasch for depositing the dCas9 plasmid on Addgene. The authors would also acknowledge Jacob Ritz for the helpful discussions as well. This work was supported by funding from the NIH R21GM27965 and Mayo Graduate School of Biomedical Sciences.

