## Supporting Information for "Characterizing Locus Specific Chromatin Structure and Dynamics with Correlative Conventional and Super Resolution imaging in living cells"

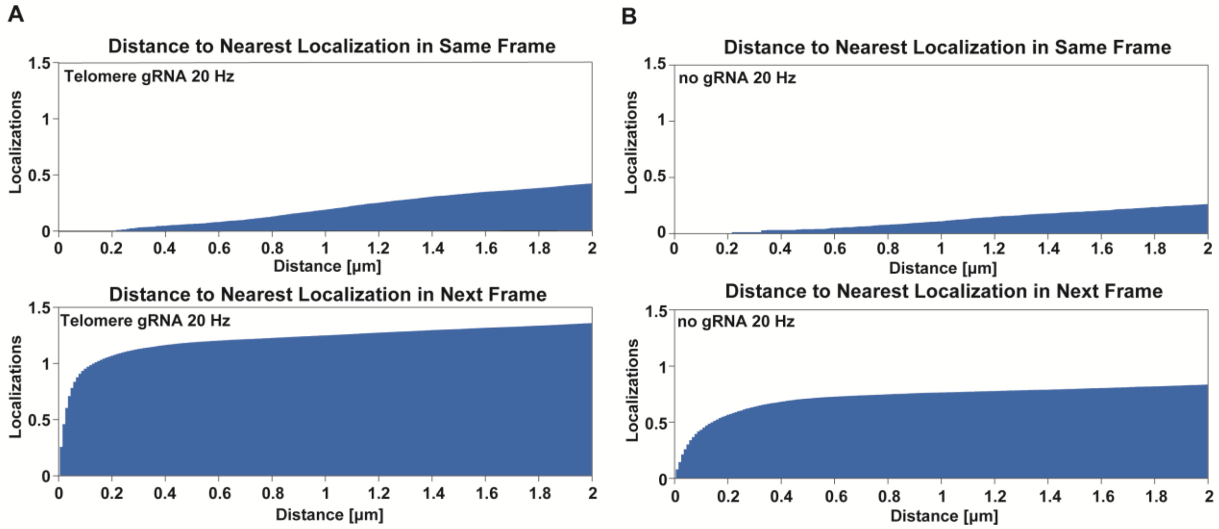


**Supplemental Figure 1: Likelihood of False Linking in Single Molecule Tracking.**

The pair correlation function quantifies how many localizations are found within a given distance to each other in the same frame (top) or next frame (bottom) **A)** For MCP-HaloTag tracking in the presence of telomere gRNA, on average two localizations are within a distance of 5.05 μm in the same frame and 120 nm in the subsequent frame. Only 0.034 localizations are within the 480 nm linking threshold in the same frame. This indicates that there is a 3.4% chance of false linking using the 480 nm linking threshold. **B)** For MCP-HaloTag tracking with no gRNA, on average two localizations are within a distance of 11.32 μm in the same frame and 3.97 μm in the subsequent frame. 0.022 localizations are found within the 480 nm linking threshold in the same frame. This indicates that there is a 2.2% chance of false linking using the 480 nm linking threshold. Data was collected from N = 5 telomere gRNA at 20 Hz and N = 5 no gRNA cells at 20 Hz.

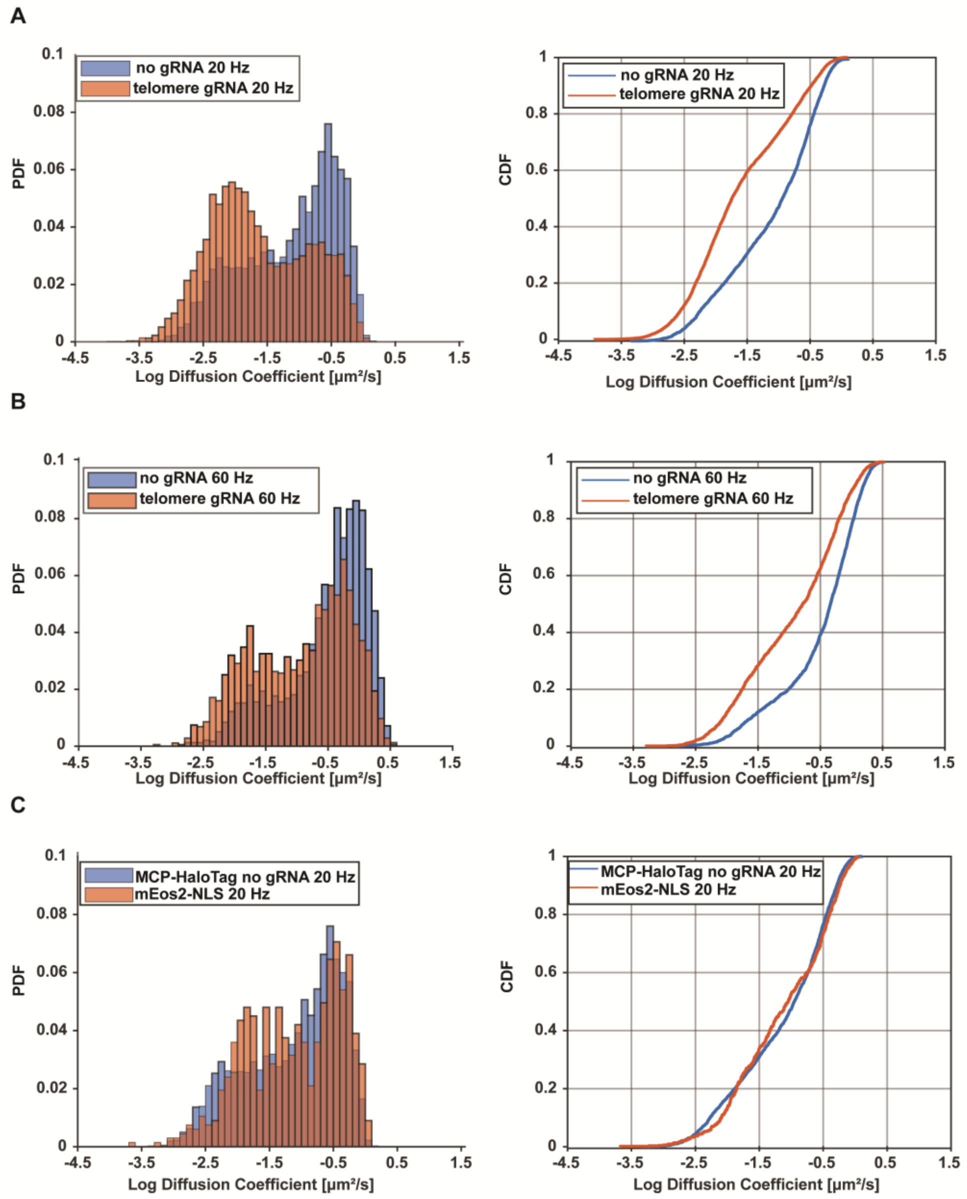


**Supplemental Figure 2: Comparison of diffusion coefficient distributions of data recorded at 20 Hz and 60 Hz frame rate. A)** Probability density function (left) and cumulative probability density function (right) of the diffusion coefficient distribution of MCP-HaloTag with and without telomere gRNA recorded at 20 Hz frame rate (same data as presented in Figure 1C). **B)** Probability density function (left) and cumulative probability density function (right) of the diffusion coefficient distribution of MCP-HaloTag with and without telomere gRNA recorded at 60 Hz frame rate (n = 4875 traces with telomere gRNA and N = 3518 traces without gRNA from N = 3 cells for each case). There was no statistically significant difference between the distributions recorded at 20 Hz and 60 Hz (Kologmorov-Smirnov Test: P = 0.65 and P = 0.72). **C)** Probability density function (left) and cumulative probability density function (right) of mEos2-NLS (N = 2936 traces from N = 3 cells) and MCP-HaloTag recorded at 20 Hz (same as MCP-HaloTag data presented in Fig 1C and Sup. Fig 2A). There was no statistically significant difference between the distributions of MCP-HaloTag and mEos2-NLS (Kologmorov-Smirnov Test P = 0.62).

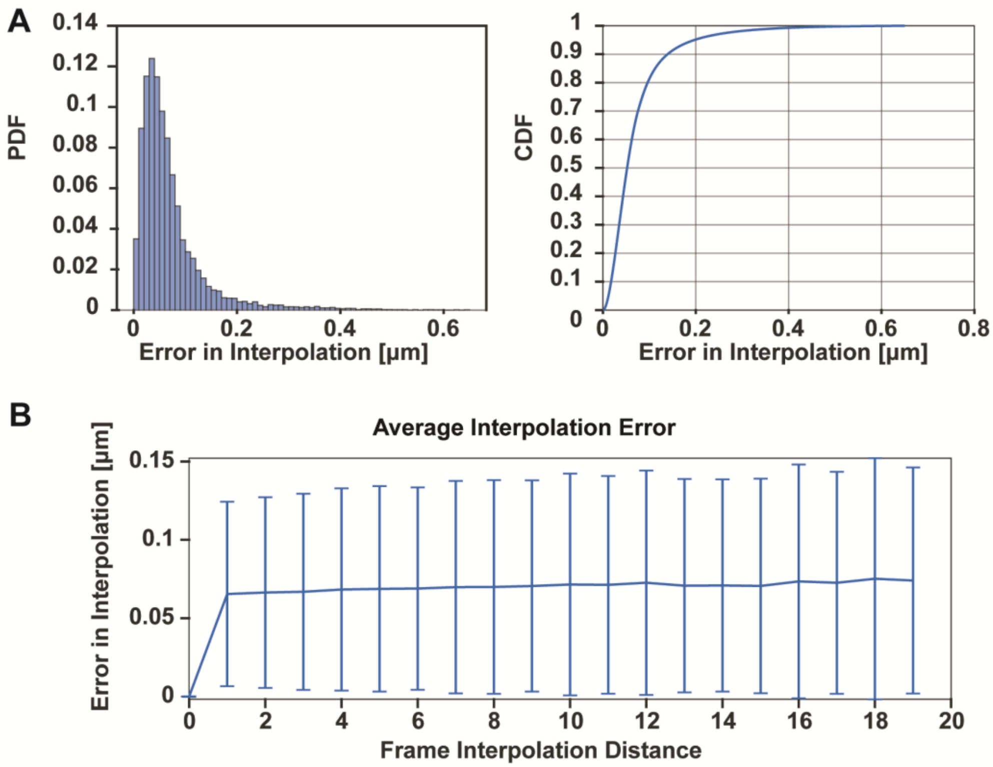


**Supplemental Figure 3: Interpolation Error of GFP Cluster Tracking**

A) Probability density function (left) and cumulative density function (right) of the interpolation error of telomere clusters. Telomeres were continuously imaged and tracked with the conventional fluorescence signal of GFP at 20 Hz for 200 seconds. To mimic the interpolation of the correlative conventional and PALM imaging data in which the telomere GFP signal is imaged localized every 10th frame, the localizations of telomeres were linearly interpolated between every 10th frame and compared to the actual localizations in the remaining frames. The distances between the centers of the interpolated positions and the actual positions were calculated to estimate the interpolation error. **B)** Average error of the interpolation across a varying number of frames. The small increase in the error if localizations are interpolated across a larger number of frames indicates that the error is not dominated by the interpolation but rather by the localization uncertainty itself. The error bars represent the standard deviation of the mean interpolation error distribution. Data represents 181 telomeres from N=5 different cells.

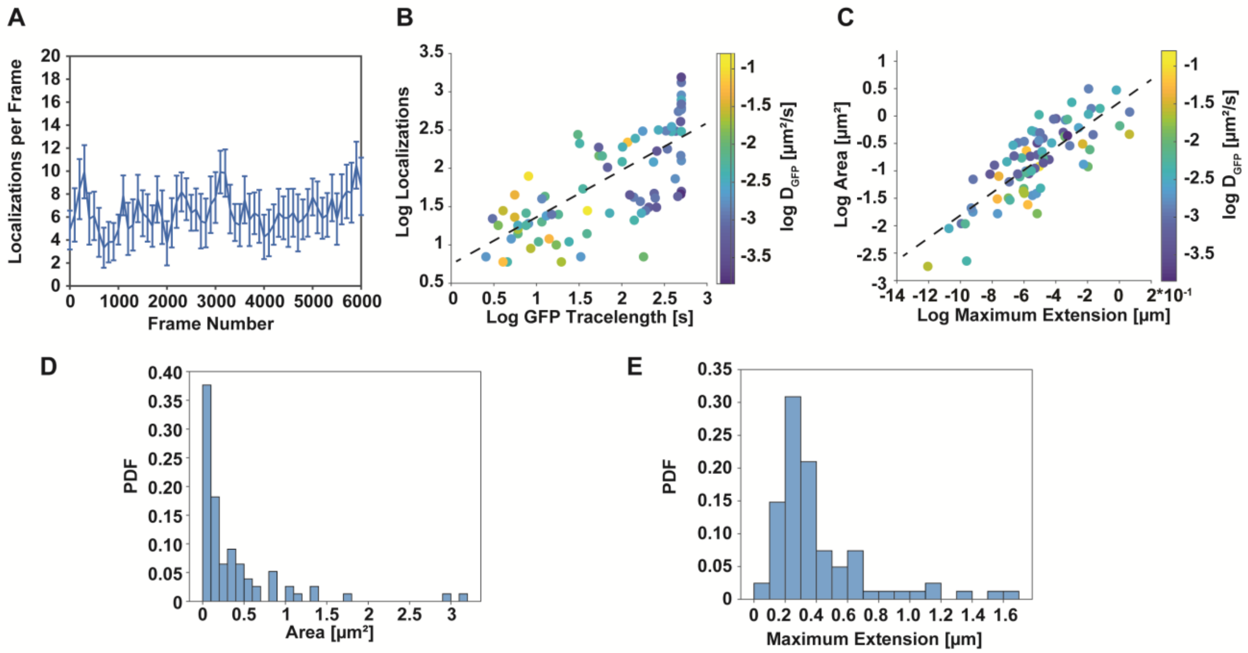

**Supplemental Figure 4: Normalization of the number of localizations by the observation time of telomeres and telomere size quantification.**

**A-B)** The detected number of PALM localizations of MCP-HaloTag per frame is constant for the duration of the movie. Therefore, the number of localizations detected for each telomere can be normalized by the time a telomere stayed in focus in order to correct for differences in the number of detected localizations. **C)** Comparison between the area of telomeres determined from motion corrected single molecule localizations and their maximum distance. While there is a correlation between the telomere area and their maximum extension (Correlation Coefficient = 0.82), deviations indicate irregular shapes. **D)** and **E)** show the histogram of telomere cluster areas and maximum extensions. All data displayed in this figure is from N = 5 cells and N = 81 telomere clusters.

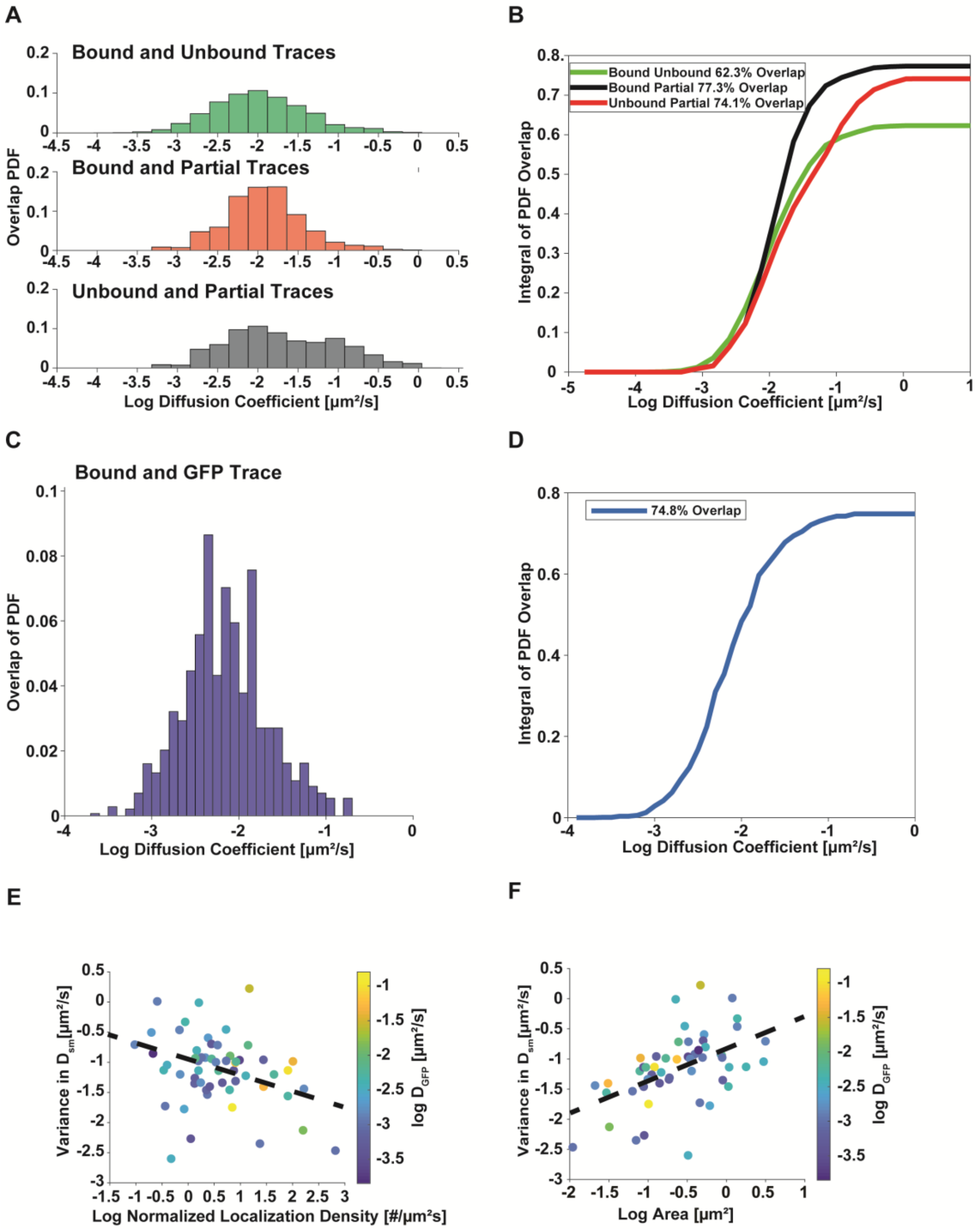


**Supplemental Figure 5: Overlap of Diffusion Coefficient Distributions Calculation and Variance in the diffusion coefficients of bound MCP-HaloTag proteins.**

**A)** Overlap integral of in the PDF of diffusion coefficient distributions shown in Figure 3A. **B)** The integral of the overlap shows the total overlap fraction as a function of the diffusion coefficient and assesses the misclassification error in when assigning a trace to a mobility state using a fixed mobility threshold. **C)** and **D)** compare the overlap in the diffusion coefficient distributions of motion corrected MCP-HaloTag traces bound to telomeres and GFP telomere cluster traces. This plot shows that there is a high percent overlap between the two distributions but differences in the both distributions remain. E) The variance of the diffusion coefficients of MCP-HaloTag proteins bound to individual telomeres shows a slight negative correlation (Correlation Coefficient = - 0.34) with the number of detected localizations. This indicates that dense telomeres exhibit less relative motion of bound MCP-HaloTag proteins. F) The variance of the diffusion coefficients of MCP-HaloTag proteins bound to individual telomeres shows a positive correlation with the area of telomeres (Correlation Coefficient = 0.47). This indicates that larger telomeres exhibit more relative motion of bound MCP-HaloTag proteins. There was no significance difference in single molecule variance across N = 5 cells as assessed by multi-way ANOVA (P = 0.25).

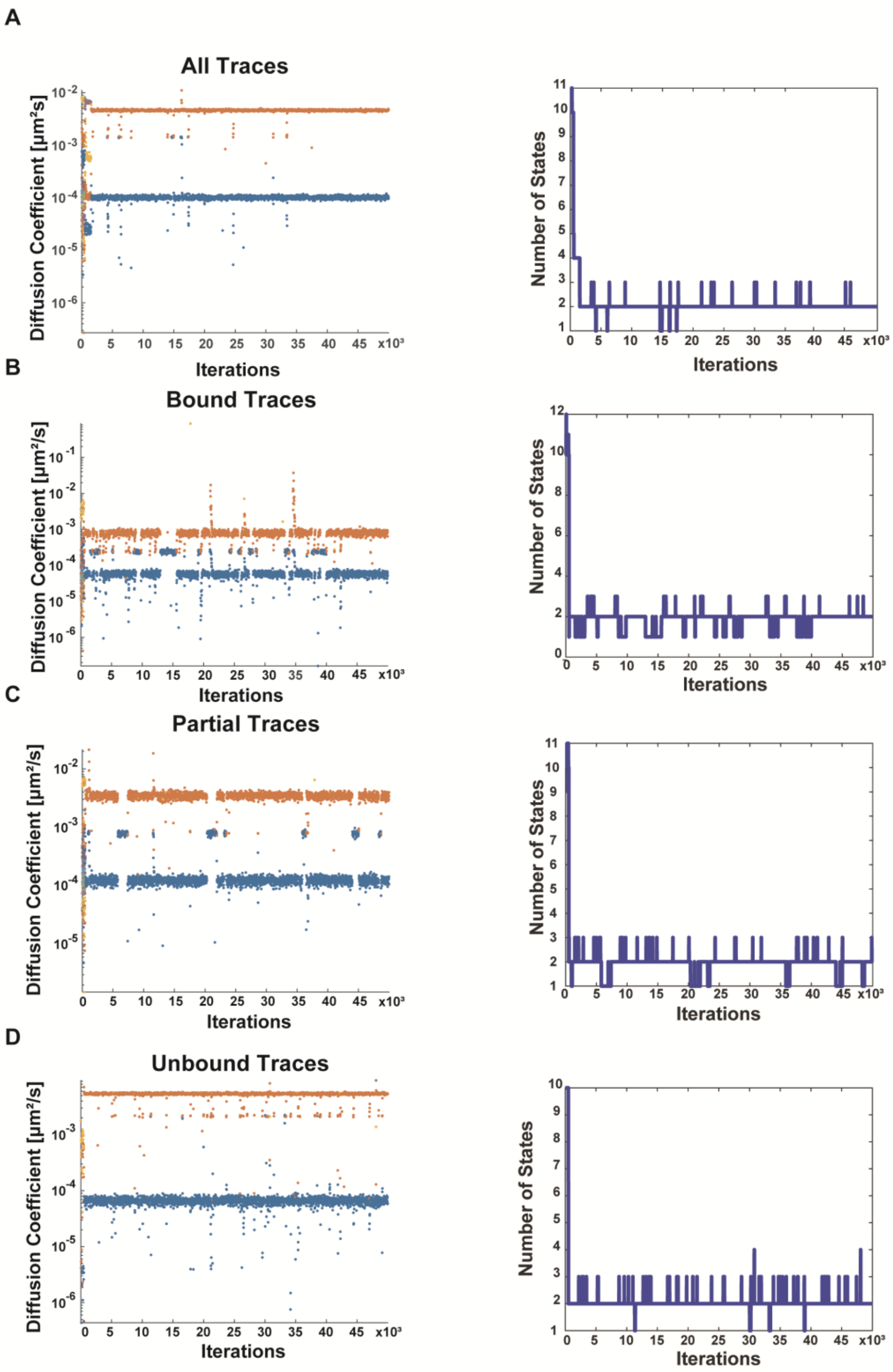


**Supplemental Figure 6: SMAUG algorithm convergence analysis**

**A)-D) Left:** Diffusion coefficients of mobility states per iteration for all traces, bound traces identified by motion correction, unbound traces identified by motion correction, and partial traces identified by motion correction. **Right**: Number of the mobility states of the SMAUG algorithm identified for each of the 50000 iterations. These plots highlight that given enough iterations, the

SMAUG algorithm converges to the same number of mobility states and similar percent of trace displacements in each mobility state.
